# Elevated blood creatinine -a biomarker of renal function-associates with multiple metabolic perturbations in dogs

**DOI:** 10.1101/2020.05.06.078063

**Authors:** Claudia Ottka, Katariina Vapalahti, Ann-Marie Määttä, Nanna Huuskonen, Sinikka Sarpanen, Liisa Jalkanen, Hannes Lohi

## Abstract

**BACKGROUND:** Renal diseases, such as chronic kidney disease (CKD) are common in dogs. While the kidneys have multiple important metabolic functions, the occurrence of metabolic disturbances in canine renal diseases has not been extensively studied.

**OBJECTIVES:** To identify metabolic changes in blood samples exhibiting elevated blood creatinine, indicating reduced renal filtration.

**ANIMALS:** Samples consisted of clinical samples analysed by a 1H NMR-based metabolomics platform. The case group included 23 samples with creatinine > 125 μmol/l, and the control group 873 samples with creatinine within the reference interval.

**METHODS:** Biomarker association with elevated creatinine was evaluated utilizing three statistical approaches: Wilcoxon rank-sum test and logistic regression analysis (FDR-corrected p-values), and classification using random forest. Means of the biomarkers were compared to reference intervals. A heatmap and histograms visualized the differences.

**RESULTS:** The levels of citrate, tyrosine, branched-chain amino acids, valine, leucine, albumin, linoleic acid % and the ratio of phenylalanine to tyrosine differed significantly both in the Wilcoxon test and logistic regression, acetate levels only in Wilcoxon test and docosapentaenoic acid % only in logistic regression (p <. 05). The ten most significant markers in random forest corresponded to the Wilcoxon test, supplemented with alanine.

**CONCLUSIONS AND CLINICAL IMPORTANCE:** This study identified multiple metabolic changes associated with elevated blood creatinine, including prospective diagnostic markers and therapeutic targets. The NMR metabolomics test is a promising tool for improving diagnostics and management of canine renal diseases. Further research is needed to verify the association of these changes to the canine patient’s clinical state.

## Introduction

Renal diseases are common in dogs, and are frequent causes of death. Renal disease is either viewed as acute or chronic, although these two can occur simultaneously and both increase the susceptibility to the other. Chronic kidney disease (CKD) is characterized by gradual renal damage, whereas acute kidney injury (AKI) is characterized by an abrupt decline in renal function. Renal damage in AKI can be reversible, whereas renal damage in CKD is typically irreversible. CKD is considered the most common renal disease in dogs, with an estimated prevalence of 0,37%^1^.

The blood creatinine concentration is a clinical biochemistry measure commonly used as a determinant of renal glomerular filtration rate. The International Renal Interest Society (IRIS) has created widely used guidelines both for the staging and treatment of CKD, and grading of AKI^2–4^. In these guidelines, grading of AKI is based on glomerular filtration capacity evaluated using the blood creatinine concentration and the presence of other clinical evidence of AKI, with further classification based on urine formation and need of renal replacement therapy^2^. The staging of CKD is based on the glomerular filtration capacity measured as blood creatinine or symmetric dimethylarginine, with further classification based on proteinuria and blood pressure^4^. Treatment guidelines are based on CKD staging, clinical signs, and plasma phosphate concentrations, as well as on possible complications; metabolic acidosis, anemia, and dehydration^3^.

However, the metabolic functions of the kidneys range further than this. Multiple additional metabolic derangements, such as impaired renal enzymatic activity^5^, have been found to occur in both human and animal renal diseases. Some of these changes have even been proposed as therapeutic targets^6–9^. However, the occurrence and significance of changes in systemic metabolism have not yet been extensively studied in dogs. Moreover, traditional diagnostic approaches do not detect the majority of these metabolic changes, hindering their use in clinical practice. We have recently developed and validated a clinically usable NMR metabolomics testing platform for dogs^10^, offering the technology for a holistic approach to both veterinary research and clinical practice.

The objective of this study was to evaluate the systemic metabolic changes occurring in dogs with elevated blood creatinine concentrations, indicating reduced renal filtration, and to discuss the possibility of these changes to be used as diagnostic biomarkers and therapeutic targets of renal diseases.

## Materials and methods

### Samples

The workflow for the study is summarized and presented in Fig 1. The study was performed as a retrospective review of clinical blood samples. The samples originated from two sources, first:

**Figure 1.**
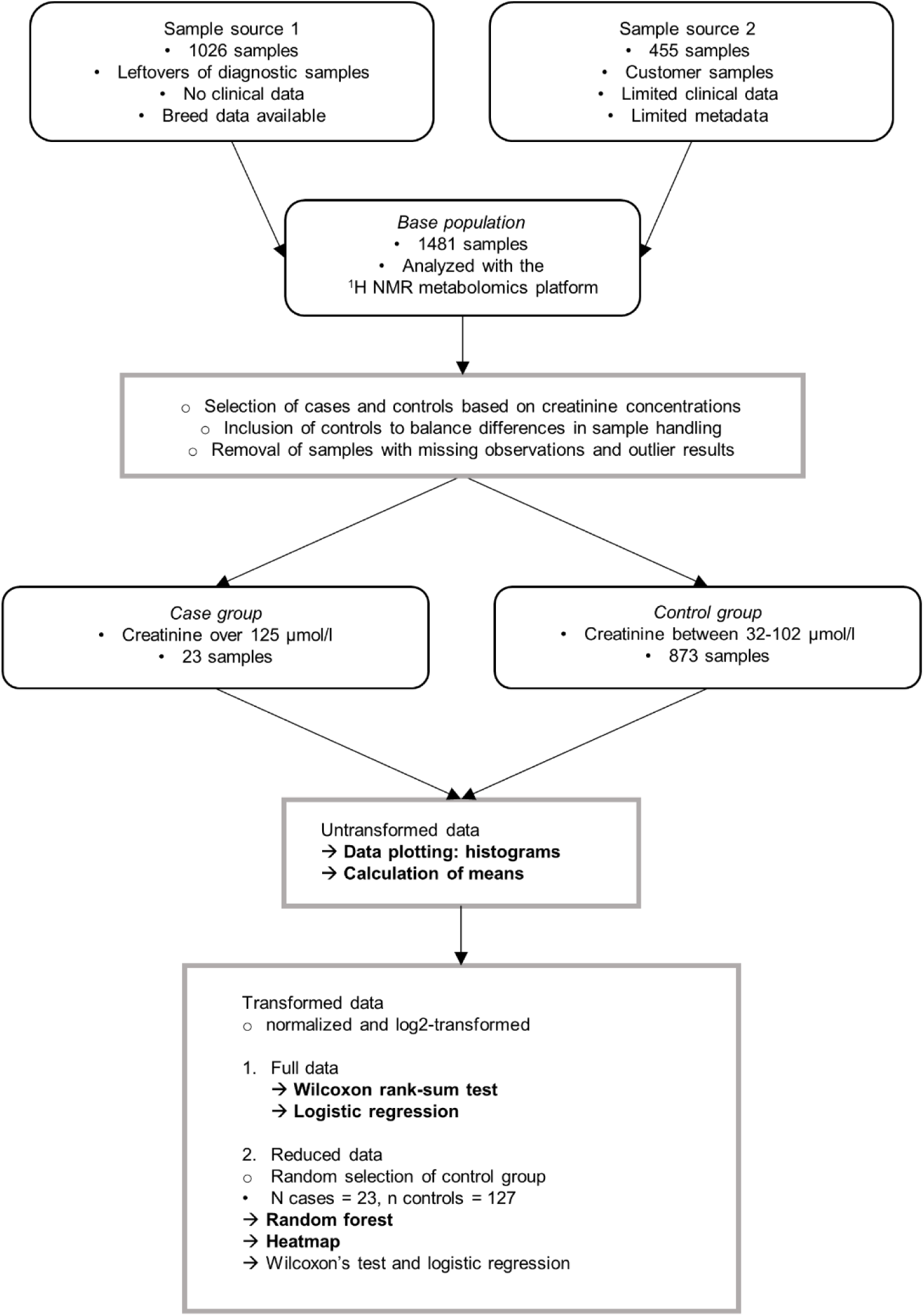
Study workflow. Rounded boxes include information on materials, boxes information on methods. Black points represent sample characteristics and circles represent data handling procedures. Arrows represent statistical analyses, and analyses primarily evaluated in this article are in bold.

Diagnostic sample material (n = 1026) taken by Finnish veterinarians, submitted by mail to a single laboratory provider (Movet Ltd., Kuopio, Finland). Altogether 999 of these samples were collected and sent during 2/2018 - 5/2018, and previously used in the method validation of the canine NMR metabolomics platform^10^, and 27 were collected and sent during 10/2018 - 2/2019 together with the samples from source 2 as described below. No clinical data was available for these samples. Signalment data was either fully missing or limited to the breed definition. Both heparin plasma and serum samples were included, and the individual sample type of each sample was unknown.

The second source included clinical samples (n = 455) collected at two Veterinary clinics (Kuopion Eläinlääkärikeskus Ltd., Kuopio, Finland and Pieneläinvastaanotto Punaturkki Ltd., Kuopio, Finland) during the NMR metabolomics clinic pilot. Samples were collected during routine veterinary appointments, and sent to NMR analysis between 10/2018 - 2/2019. Limited disease data and signalment data were available for these samples. All samples were heparin plasma samples.

The samples were analyzed by a validated canine ^1^H NMR metabolomics platform10 quantifying 123 biomarkers.

Case and control groups were created according to their NMR-measured creatinine concentrations. To limit possible confounding factors, clinical data was reviewed for samples, for which it was available. No samples were excluded based on clinical data. Since CKD is the most common renal disease in dogs, we used creatinine concentrations from IRIS CKD staging guidelines as inclusion criteria to the case group. Inclusion in the case group was based on the creatinine measurement being above 125 µmol/l, indicating CKD stage 2 or higher^4^.

The first inclusion criteria for the control group was the NMR-measured creatinine concentration being within the creatinine reference interval (32 - 103 µmol/l) of the NMR-method^10^. To minimize the confounding effect of preanalytical variability caused by sample handling, our second inclusion criteria was to include control group samples from different sample sets in a similar ratio as cases.

### Ethical Approval

The study was performed as a retrospective evaluation of clinical blood samples and leftovers of clinical samples. All applicable international, national, and/or institutional guidelines for the care and use of animals were followed. All procedures performed in studies involving animals were in accordance with the ethical standards of the institution or practice at which the studies were conducted. Committee: Finnish national Animal Experiment Board, permit number: ESAVI/7482/04.10.07/2015. Permission for scientific use was obtained for all samples.

### Statistical analysis

The data was evaluated for missing observations. We removed biomarkers with multiple missing observations and samples with missing observations. Two samples in the case group showed marked lipemia with the measured triglyceride concentration of over 7 mmol/l. In order to remove skewness caused by these lipemic samples, we removed these samples from further analyses as potential outliers. The resulting case group consisted of 23 samples and control group of 873 samples.

We used three different statistical approaches to identify metabolite association with elevated creatinine concentrations. Before conducting statistical analyses, the data was normalized and log2-transformed to reduce bias and to balance the data. The three statistical approaches included

i. Wilcoxon rank-sum test with FDR-corrected p-values to evaluate the significance of differences between case and control groups’ analyte concentrations,
ii. separate logistic regression models for each biomarker to evaluate the metabolite’s association to elevated creatinine concentrations. The case-control status served as the response variable and the individual metabolite as the independent variable. To control the Type I errors in multiple testing, we used FDR p-value correction. Linearity between the metabolites and log odds were tested. The goodness of fit and predictive value of the models were assessed by AIC and AUC values – a lower AIC indicates better fit and a higher AUC a better predictive value.
iii. random forest classification to identify the biomarkers, that predicted best the elevated creatinine concentrations. To balance the difference in number of cases and controls, control group size was reduced by random undersampling to 127 samples. Biomarker association with elevated creatinine concentrations was assessed by variable importance.

To check, whether metabolite association with elevated creatinine concentrations is the same using the reduced and full control groups, we performed the Wilcoxon rank-sum test and logistic regression analysis also with the reduced control group.

A heatmap was created to visualize the results of the biomarkers associated with elevated creatinine concentrations based on the Wilcoxon rank-sum test, logistic regression and random forest, using the reduced data of 127 samples.

To determine the direction of the observed changes and to evaluate, whether the changes would be detected in clinical diagnostics based on reference intervals, we compared the means of the case and control groups to each other and to analysis reference intervals. We evaluated, whether the means of untransformed metabolite values and their 95 % confidence intervals (CI) in the case group markedly differ from the reference intervals of the NMR method8, and their 90% CI.

To visualize the analyte concentrations in the case and control group compared to reference intervals of the NMR method, we plotted histograms of the analytes. Potential outliers were removed from the control group to ensure optimal histogram scaling.

All statistical analyses conducted throughout this study were performed by SAS version 9.4, SAS Institute Inc., Cary, NC, USA, RStudio Team (2019). RStudio: Integrated Development for R. RStudio, Inc., Boston, MA URL http://www.rstudio.com/and Microsoft Office Excel, Microsoft Corp., Redmond, WA, US.

## Results

### Sample characteristics

In this study, we used a non-targeted ^1^H NMR metabolomics approach to compare metabolite profiles of clinical samples with elevated creatinine concentrations (n = 23) to clinical samples with normal creatinine concentrations (n = 873). The median creatinine concentration in the case group was 196 µmol/l, with a range of 131 - 646 µmol/l. The median creatinine concentration in the control group was 60 µmol/l with a range of 32 - 102 µmol/l. Since most of the used samples were leftovers of clinical laboratory samples, limited signalment and clinical data was available. Breed was unknown for 10 (43.5 %) of the case group and 186 (21.3 %) of the control group samples (Table 1). The remaining 13 samples in the case group were taken from dogs of 12 different breeds. The control group samples were taken from dogs of 155 different breeds, including also 8 % of mixed breed dogs.

**Table 1.** Summary of the most common breeds in the case and control groups.

| Cases |  | Controls |  |
| --- | --- | --- | --- |
| Breed | % | Breed | % |
| Unknown | 43.5 | Unknown | 21.3 |
| Lagotto Romagnolo | 8.7 | Mixed breed | 8.0 |
| Poodle | 4.3 | Labrador Retriever | 3.8 |
| Nova Scotia Duck Tolling Retriever | 4.3 | German Shepherd Dog | 2.7 |
| Bichon Frise | 4.3 | Golden Retriever | 2.4 |
| Bernese Mountain Dog | 4.3 | Shetland Sheepdog | 2.2 |
| Schapendoes | 4.3 | Finnish Lapphund | 2.2 |
| Collie Rough | 4.3 | Poodle | 1.9 |
| Miniature Pinscher | 4.3 | Spanish Water Dog | 1.8 |
| Cavalier King Charles Spaniel | 4.3 | Jack Russell Terrier | 1.6 |
| American Staffordshire Terrier | 4.3 | Nova Scotia Duck Tolling Retriever | 1.5 |
| Dogue de Bordeaux | 4.3 | Swedish Elkhound | 1.3 |
| Bouvier des Flandres | 4.3 | Finnish Hound | 1.3 |
|  |  | Miniature Schnauzer | 1.1 |
|  |  | Bichon Frise | 1.1 |

The table shown 15 most common breeds in the case and control groups, including samples with unknown breed and mixed breeds. The total number of case group samples was 23, and control group samples was 873. The total number of breeds in the control group was 155, including mixed breed dogs, and 45,8 % of the control group samples were taken from breeds not included in the 15 most common breeds in the control group.

The used NMR metabolomics platform quantitates 123 biomarkers. Biomarkers with multiple missing observations were excluded from further analyses, resulting in 97 analyzed biomarkers (Supplementary Table 1).

### Results of the statistical association tests

The Wilcoxon rank-sum test, logistic regression analysis and classification using random forest identified similar biomarkers (Table 2, Fig 2).

**Table 2.**
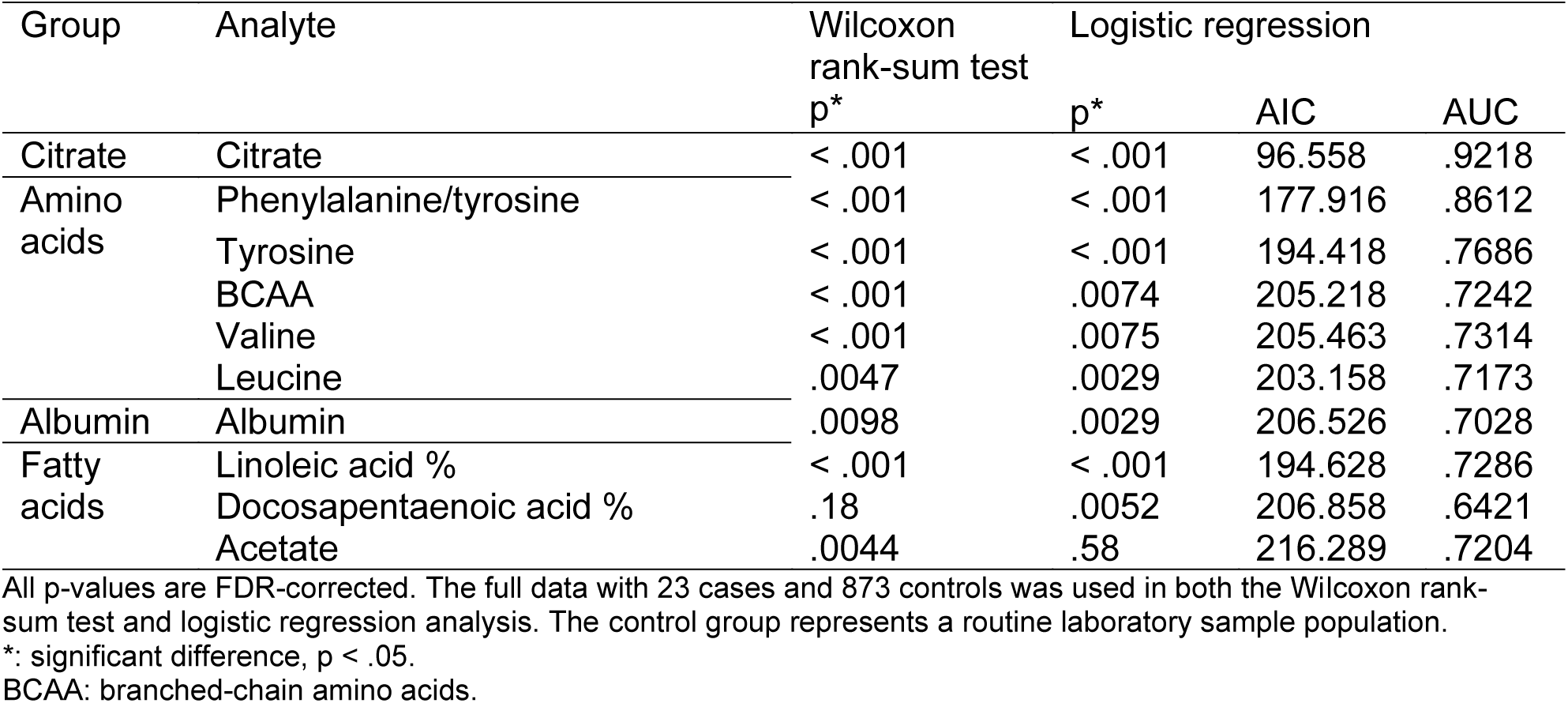
Analytes with significant differences between the case and control group according to Wilcoxon rank-sum test and logistic regression analysis.

| Group | Analyte | Wilcoxon rank-sum test | Logistic regression |  |  |
| --- | --- | --- | --- | --- | --- |
|  |  | p* | p* | AIC | AUC |
| Citrate | Citrate | < .001 | < .001 | 96.558 | .9218 |
| Amino acids | Phenylalanine/tyrosine | < .001 | < .001 | 177.916 | .8612 |
|  | Tyrosine | < .001 | < .001 | 194.418 | .7686 |
|  | BCAA | < .001 | .0074 | 205.218 | .7242 |
|  | Valine | < .001 | .0075 | 205.463 | .7314 |
|  | Leucine | .0047 | .0029 | 203.158 | .7173 |
| Albumin | Albumin | .0098 | .0029 | 206.526 | .7028 |
| Fatty acids | Linoleic acid % | < .001 | < .001 | 194.628 | .7286 |
|  | Docosapentaenoic acid % | .18 | .0052 | 206.858 | .6421 |
|  | Acetate | .0044 | .58 | 216.289 | .7204 |

All p-values are FDR-corrected. The full data with 23 cases and 873 controls was used in both the Wilcoxon rank-sum test and logistic regression analysis. The control group represents a routine laboratory sample population.
\*: significant difference, $p < .05$ .
BCAA: branched-chain amino acids.

**Figure 2.**
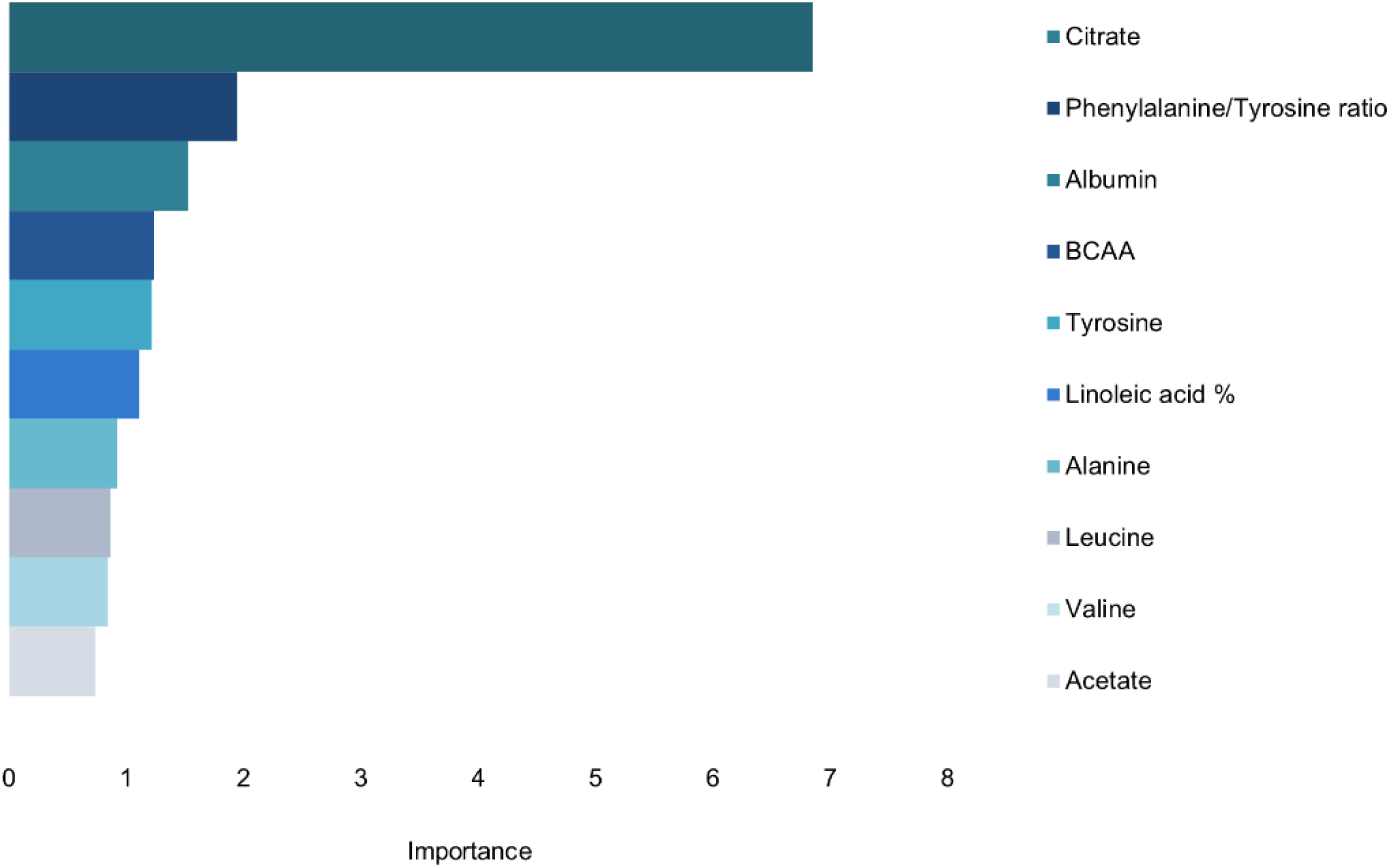
Variable importance in random forest classification using the reduced data. Ten biomarkers with the highest variable importance are included in the figure. The higher the variable importance, the more important the feature is in predicting high creatinine concentration. Random forest classification was conducted using the reduced data with 23 cases and 127 controls. The control group represents a routine laboratory sample population. BCAA: branched-chain amino acids.

Citrate, tyrosine, branched-chain amino acids (BCAA), valine, leucine, albumin, acetate, linoleic acid % and the ratio of phenylalanine to tyrosine showed significant differences between cases and controls in the Wilcoxon rank-sum test (Table 2). The same biomarkers, excluding acetate, and including docosapentaenoic acid % were associated with elevated creatinine concentrations in logistic regression analysis. The best model fit and predictive values (AIC and AUC, respectively) were achieved for citrate, followed by phenylalnine/tyrosine and tyrosine.

In classification using random forest, the ten biomarkers with the highest variable importance were the same biomarkers that reached significance in the Wilcoxon rank-sum test; citrate, tyrosine, BCAA, valine, leucine, albumin, acetate, linoleic acid % and the ratio of phenylalanine to tyrosine, as well as the amino acid alanine (Fig 2).

Since we used different sample sets for different statistical approaches; the full data set for logistic regression analysis and Wilcoxon rank-sum test, and a reduced data set with randomly selected samples for random forest classification, we checked whether results of logistic regression and the Wilcoxon rank-sum test are the same in the full and reduced data sets. Both data sets gave similar results.

The heatmap visualized the aforementioned differences in metabolite levels in case and control samples, but it also showed large amounts of variability within both the case and control group samples (Fig 3).

**Figure 3.**
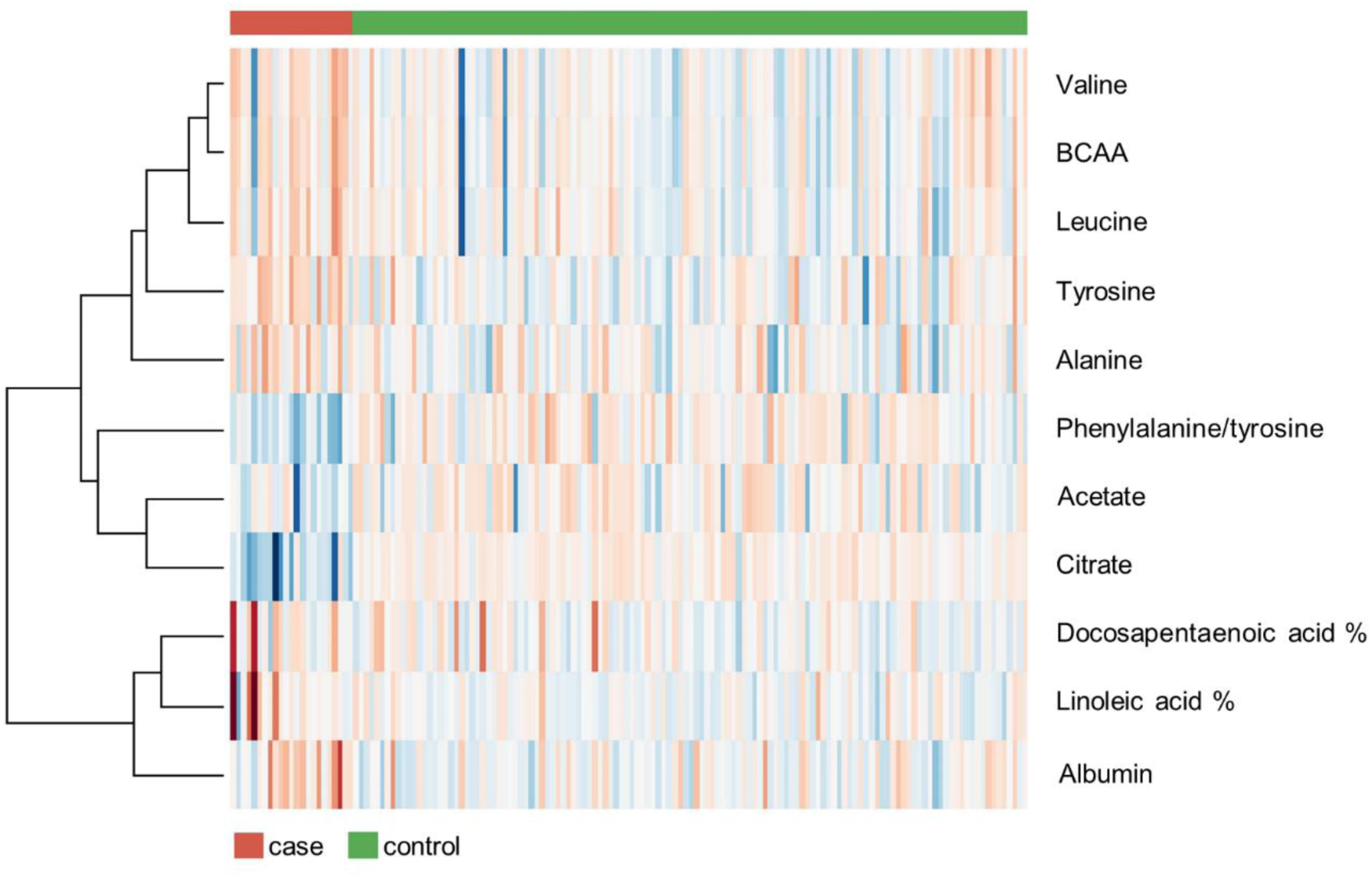
Heatmap of the reduced data. Metabolites associated with elevated creatinine concentrations according to the Wilcoxon rank-sum test, logistic regression analysis or random forest classification are included. The heatmap was created using the reduced data with 23 cases and 127 controls, with normalized and log2-transformed data. The control group represents a routine laboratory sample population. Each column represents one sample. Columns with a red line on top represent the case group, and the columns with a green line on top represent the control group. In the subsequent rows, red hues represent elevated levels and blue hues decreased levels. Color intensity increases proportionally to the magnitude of the change.

### Magnitudes of the differences in analyte concentrations: means and histograms

The concentrations of BCAA, leucine, valine, tyrosine, alanine, albumin, docosapentaenoic acid % and linoleic acid % were lower in the case than control group, whereas mean acetate, citrate and phenylalanine/tyrosine ratio were higher in the case group (Table 3). The case group’s mean citrate concentration and phenylalanine/tyrosine ratio were higher than the reference interval of the NMR method, indicating that the mean concentration in the case group is considered a clinically observable change.

**Table 3.** Mean concentrations of biomarkers associated with elevated creatinine concentrations according to the Wilcoxon rank-sum test, logistic regression analysis or random forest classification.

| Analyte | Mean case ±<br>CI | Mean control ±<br>CI | RI (CI)<sup>10</sup> |
| --- | --- | --- | --- |
| Citrate mmol/l | .156 ± .027<sup>a</sup> | .077 ± .001 | .061 (.059 - .063) - .123 (.120 - .125) |
| Phenylalanine/tyrosine<br>ratio | 1.114 ± .103<sup>b</sup> | .794 ± .013 | .513 (.494 - .530) - 1.049 (1.026 -<br>1.098) |
| Tyrosine mmol/l | .052 ± .005 | .066 ± .001 | .041 (.039 - .043) - .089 (.086 - .091) |
| BCAA mmol/l | .334 ± .044 | .394 ± .006 | .242 (.215 - .251) - .515 (.502 - .531) |
| Valine mmol/l | .165 ± .026 | .195 ± .003 | .113 (.099 - .119) - .251 (.243 - .261) |
| Leucine mmol/l | .109 ± .013 | .132 ± .002 | .083 (.078 - .088) - .185(.181 - .189) |
| Alanine mmol/l | .356 ± .043 | .407 ± .007 | .216 (.199 - .222) - .597 (.580 - .624) |
| Albumin g/l | 27.0 ± 1.1 | 28.8 ± .1 | 25.2 (24.2 - 25.7) - 32.4 (32.1 - 32.8) |
| Linoleic acid % | 25.6 ± 1.2 | 27.1 ± .1 | 24.1 (23.9 - 24.4) - 28.6 (28.4 - 28.7) |
| Docosapentaenoic acid % | 1.1 ± .0 | 1.3 ± .0 | 1.1 (1.0 - 1.1) - 2.0 (1.9 - 2.0) |
| Acetate mmol/l | .031 ± .004 | .026 ± .001 | .021 (.020 - .021) - .037 (.036 - .037) |

The full data with 23 cases and 873 controls was used when calculating case and control group means. The control group represents a routine laboratory sample population. Reference intervals represent the reference intervals of the NMR method<sup>10</sup>.
<sup>a</sup> Analytes mean and 95% CI in the case group higher than the analysis reference interval and its 90% CI.
<sup>b</sup> Analytes mean in the case group higher than the analysis reference interval; overlap in analytes 95% CI and reference intervals 90% CI.

Histograms of the analytes confirmed, that citrate levels and phenylalanine/tyrosine ratio are notably higher in samples with elevated creatinine, and often exceed the reference interval (Fig 4). The dispersion of citrate was high in the case group. The histogram of linoleic acid % showed multiple outlier results in the case group, and control group results approached the upper reference limit of the analysis. The histogram of acetate suggested, that control group results approached the lower analysis reference limit. Although the distribution of albumin approached the lower reference limit, certain samples had their albumin concentration near the upper reference limit.

**Figure 4.**
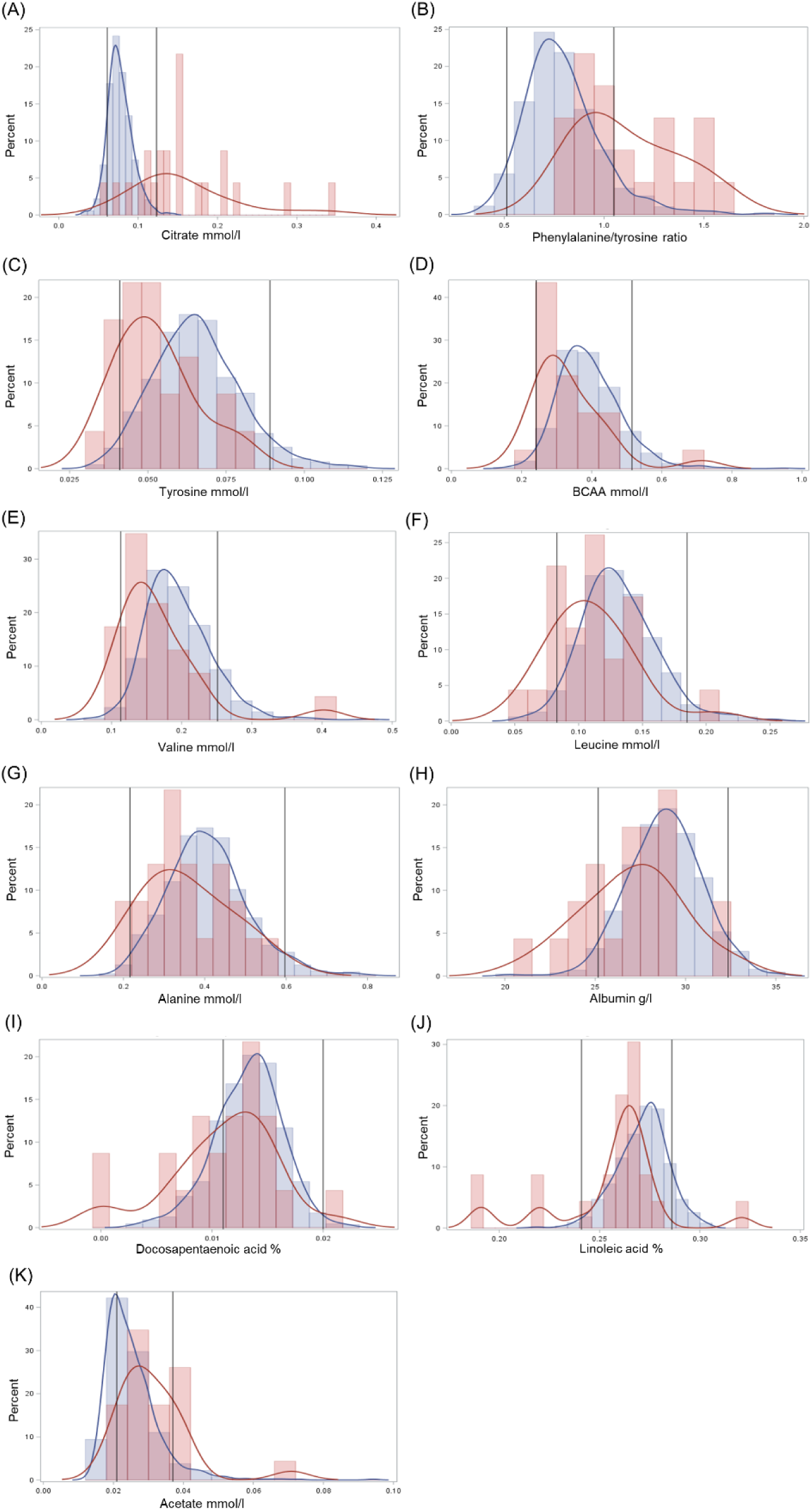
A-K: Histograms of biomarkers associated with elevated creatinine concentration according to the Wilcoxon rank-sum test, logistic regression analysis or random forest classification. The full, untransformed data with 23 cases and 873 controls was used to create histograms. The control group represents a routine laboratory sample population. Red lines and bars: case group. Blue lines and bars: control group. Black lines: reference intervals of the NMR method10. BCAA: Branched-chain amino acids.

## Discussion

Impaired renal function is common in dogs, and CKD is a common cause of death among senior dogs. Blood creatinine measurements are routinely used as a measure of renal function, with elevated creatinine indicating reduced renal filtration. Systemic metabolic derangements occurring during impaired renal function have gained attention recently, but have not yet been extensively studied in dogs, and are therefore not yet utilized in clinical practice. The aim of this study was to utilize our novel quantitative metabolomics test to identify circulating metabolites associated with the elevated blood creatinine concentration, and to evaluate their usability in renal disease management. We discovered a broad-range of physiologically relevant metabolites, such as citrate, branched-chain amino acids, albumin and fatty acids associated to reduced renal filtration. These findings may give new insights to compromised renal function and opportunities for the clinical management of renal diseases.

Blood citrate levels were markedly elevated in cases, and showed high dispersion of the results, suggesting different levels of altered citrate metabolism in these animals. Elevated blood citrate concentrations have previously been associated with impaired renal function in both humans and rats^11,12^. The kidneys are responsible for citrate removal from plasma by both urinary excretion, and renal tubular cell citrate metabolism, which contributes to renal energy supply^13^. An important determinant of citrate excretion is blood pH, and increased citrate reabsorption can occur due to acid retention and metabolic acidosis^7,14^. The use of dietary H+ reduction has been suggested in CKD patients with reduced citrate excretion^7^.

The aromatic amino acids phenylalanine and tyrosine have been associated with CKD^5,9,15^. In normal conditions, tyrosine is considered a non-essential amino acid, since the body is capable of forming it sufficiently from phenylalanine. In CKD, however, phenylalanine hydroxylation to tyrosine can be reduced^5,15^. Insufficient phenylalanine hydroxylation to tyrosine leads to a decrease in plasma tyrosine concentrations and a normal to slightly increased plasma concentration of phenylalanine, and an increase in the plasma ratio of phenylalanine to tyrosine^5,9,16^. These changes were also observed in this study, with the tyrosine concentration being significantly lower, and the ratio of phenylalanine to tyrosine being significantly higher in the case than the control group. It has been suggested, that dietary tyrosine supplementation should be used when renal tyrosine formation is insufficient^6,9^. Further studies are needed to identify the tyrosine concentrations, which would benefit from tyrosine supplementation.

We observed also significantly lower levels of total branched-chain amino acids (BCAA) as well as the individual BCAAs valine and leucine in dogs with elevated creatinine compared to the routine laboratory population. The concentrations of BCAA are known to fall in CKD^9,17^, especially during metabolic acidosis^18,19^. This phenomenon is caused by increased catabolism of muscle and branched chain amino acids due to the increased activity of liver and muscle branched-chain keto acid dehydrogenase, with reduced protein intake also contributing to the condition^20^. This condition can be treated in human medicine by supplementing the low-protein diet with BCAA or their keto analogues (BCKA)^21,22^. BCAA/BCKA supplementation has also been suggested for hypoalbuminemic and hypoaminoacidemic dogs, whose condition is not adequately controlled by routine treatments including clinical renal diets and ACE-inhibitors^8^.

Observing both hypo- and hyperalbuminemia in the case group suggests, that both chronic and acute renal pathologies were included in the case group. Hypoalbuminemia is most commonly observed in CKD and caused by urinary leakage of albumin. Since albumin is responsible for the maintenance of blood oncotic pressure, severe hypoalbuminemia causes fluid leakage out of the blood vessels, causing edema or ascites. Proteinuria is one of the assessed parameters in the IRIS CKD guidelines, and affects treatment of the disease^3,4^. Hyperalbuminemia, on the other hand, can occur due to polyuria and vomiting, especially when water is withheld, and is a common feature of AKI. Correction of fluid balance is a vital treatment goal in these patients, since dehydration can cause ischemic kidney injury.

The acetate concentration was significantly higher in case samples than in control samples. Elevated blood acetate concentrations have previously been observed in a rat model of CKD^12^. Short-chain fatty acids (SCFA), such as acetate, are organic anions that are largely produced by gut microbial fermentation^23^. SCFA have immunomodulatory effects^24^ and affect renal blood pressure regulation^25^. Long-term administration of large doses of SCFA has been associated with the development of ureteritis and hydronephrosis^26^. However, since the control group acetate concentrations were relatively low in this study, further studies of acetate concentrations in canine renal disease are warranted.

The omega-3 fatty acid docosapentaenoic acid % was significantly lower in case group samples than in controls. The omega-6 fatty acid linoleic acid % was also significantly lower in case group, but control group results were slightly skewed towards the upper reference limit of the analysis, suggesting the possibility of physiological changes in the control group consisting of a routine laboratory sample population. Omega-3 fatty acids are considered renoprotective, whereas omega-6 fatty acids are considered detrimental to renal function^27^. Due to their bioactive role, omega-3 fatty acids are supplemented in functional renal diets^28^. Further studies are needed to confirm the association of fatty acid levels with the clinical state of the patient.

Alanine concentrations were associated with elevated creatinine concentrations, with the case group having lower alanine concentrations than the control group. Previously, reduced alanine concentrations have been found in human CKD patients with impaired renal function^29,30^, and urinary alanine excretion has been associated with proteinuria^30^.

Two case group samples exhibited severe hypertriglyceridemia of over 7 mmol/l and were excluded from further data analyses due to extreme outlier results. Hypercholesterolemia and mild hypertriglyceridemia are considered components of canine nephrotic syndrome, characterized by severe proteinuria^31^. In human patients, hyperlipidemia typically occurs also in CKD patients without proteinuria, and hyperlipidemia has recently been reported in non-proteinuric dogs with CKD^32^. The development of hyperlipidemia is considered multifactorial; both reduced fat catabolism and increased hepatic lipoprotein formation to correct low oncotic pressure are suggested to contribute to this condition^33^. Monitoring of hyperlipidemia is important in severely hyperlipidemic patients, since it is associated with serious diseases, such as pancreatitis, vacuolar hepatopathy and gallbladder mucocele^34^. It has been recommended that persistent, severe hypertriglyceridemia exceeding 5,5 mmol/l should be treated to prevent these possible complications, however, complications might occur even at lower triglyceride levels^34^.

The major limitation of this study is sample inclusion based only on blood creatinine concentrations, not the presence of diagnosed renal disease. This approach was taken, since clinical data was available only for a small minority of the samples. Due to the inclusion criteria, the case group most likely includes samples with diverse, acute and chronic conditions affecting renal function. We also could not evaluate the effects of other physiological parameters, such as age and sex, since complete demographic data was lacking from most samples. Using routine clinical laboratory samples as the control group enabled us to find metabolites differentiating samples with high creatinine from other diseases. It also allowed us to use samples with similar sample handling procedures. However, general changes associating with multiple disease states are not well visualized by this approach, and variability in metabolite results within the control group was high.

In summary, we identified metabolic changes associated with elevated creatinine concentrations with possible implications on disease diagnostics and management. The quantitative NMR metabolomics approach is a promising tool for highlighting the metabolic alterations occurring in renal diseases and monitoring these changes in response to treatment. Further studies with well-defined clinical cohorts are needed to evaluate, how these changes associate with the specific clinical status of the canine patients.

## Supporting information

Supplementary table 1. Analyzed biomarkers

## Acknowledgements

We thank the staff of Movet Ltd. for sample acquisition and coordination of sample flow. The staff of Kuopion Eläinlääkärikeskus Ltd., and Pieneläinvastaanotto Punaturkki Ltd. are acknowledged for help in sample handling and acquisition. We thank the customers of Movet Ltd. for enabling the scientific use of leftover diagnostic sample material. We thank all dog owners, that participated in the metabolomics clinic pilot for enabling the scientific use of their dogs’ samples and metadata. We thank Kibble Labs Ltd. for NMR analysis of the samples and MediSapiens Ltd. for sample data management. The canine genetics research group at the University of Helsinki is thanked for help in sample management. Tuomas Poskiparta and Katja Jauni are acknowledged for advice and help in project conceptualization. We thank PetBIOMICS Ltd. and Academy of Finland (308887) for funding the study.

## References

1. O’Neill DG, Elliott J, Church DB, McGreevy PD, Thomson PC, Brodbelt DC. Chronic Kidney Disease in Dogs in UK Veterinary Practices: Prevalence, Risk Factors, and Survival. J Vet Intern Med [Internet]. 2013;27(4):814–21. Available from: https://onlinelibrary.wiley.com/doi/abs/10.1111/jvim.12090

2. International Renal Interest Sociery [IRIS]. Grading of Acute Kidney Injury (modified 2016). 2016.

3. International Renal Interest Society [IRIS]. Treatment recommendations for CKD in Dogs (modified 2019) [Internet]. 2019. Available from: http://www.iris-kidney.com/pdf/IRIS-DOG-Treatment_Recommendations_2019.pdf

4. International Renal Interest Society [IRIS]. IRIS Staging of CKD (modified 2019). 2019.

5. Young GA, Parsons FM. Impairment of phenylalanine hydroxylation in chronic renal insufficiency. Clin Sci. 1973 Jul;45(1):89–97.

6. Kopple JD. Phenylalanine and tyrosine metabolism in chronic kidney failure. J Nutr. 2007 Jun;137(6 Suppl 1):1586S–1590S; discussion 1597S–1598S.

7. Goraya N, Simoni J, Sager LN, Madias NE, Wesson DE. Urine citrate excretion as a marker of acid retention in patients with chronic kidney disease without overt metabolic acidosis. Kidney Int [Internet]. 2019 May 1;95(5):1190–6. Available from: https://doi.org/10.1016/j.kint.2018.11.033

8. Zatelli A, D’Ippolito P, Roura X, Zini E. Short-term effects of dietary supplementation with amino acids in dogs with proteinuric chronic kidney disease. Can Vet J = La Rev Vet Can. 2017 Dec;58(12):1287–93.

9. Parker VJ, Fascetti AJ, Klamer BG. Amino acid status in dogs with protein-losing nephropathy. J Vet Intern Med. 2019 Mar;33(2):680–5.

10. Ottka, C., Vapalahti, K., Puurunen, J., Vahtera, L. & Lohi, H. Characteristics of a novel NMR-based metabolomics platform for dogs. bioRxiv (2019). doi:10.1101/871285

11. Barrios C, Zierer J, Wurtz P, Haller T, Metspalu A, Gieger C, et al. Circulating metabolic biomarkers of renal function in diabetic and non-diabetic populations. Sci Rep. 2018 Oct;8(1):15249.

12. Kim J-A, Choi H-J, Kwon Y-K, Ryu DH, Kwon T-H, Hwang G-S. 1H NMR-based metabolite profiling of plasma in a rat model of chronic kidney disease. PLoS One. 2014;9(1):e85445.

13. Simpson DP. Citrate excretion: a window on renal metabolism. Am J Physiol Physiol [Internet]. 1983;244(3):F223–34. Available from: https://doi.org/10.1152/ajprenal.1983.244.3.F223

14. Brennan S, Hering-Smith K, Hamm LL. Effect of pH on citrate reabsorption in the proximal convoluted tubule. Am J Physiol. 1988 Aug;255(2 Pt 2):F301–6.

15. Wang M, Vyhmeister I, Swendseid ME, Kopple JD. Phenylalanine Hydroxylase and Tyrosine Aminotransferase Activities in Chronically Uremic Rats. J Nutr [Internet]. 1975;105(1):122–7. Available from: https://doi.org/10.1093/jn/105.1.122

16. Fukuda S, Kopple JD. Chronic uremia syndrome in dogs induced with uranyl nitrate. Nephron. 1980;25(3):139–43.

17. Holecek M, Sprongl L, Tilser I, Tichy M. Leucine and protein metabolism in rats with chronic renal insufficiency. Exp Toxicol Pathol. 2001 Apr;53(1):71–6.

18. Hara Y, May RC, Kelly RA, Mitch WE. Acidosis, not azotemia, stimulates branched-chain, amino acid catabolism in uremic rats. Kidney Int. 1987 Dec;32(6):808–14.

19. May RC, Masud T, Logue B, Bailey J, England BK. Metabolic acidosis accelerates whole body protein degradation and leucine oxidation by a glucocorticoid-dependent mechanism. Miner Electrolyte Metab. 1992;18(2-5):245–9.

20. Holecek M. Branched-chain amino acids in health and disease: metabolism, alterations in blood plasma, and as supplements. Nutr Metab (Lond). 2018;15:33.

21. Cupisti A, Brunori G, Di Iorio BR, D’Alessandro C, Pasticci F, Cosola C, et al. Nutritional treatment of advanced CKD: twenty consensus statements. J Nephrol. 2018 Aug;31(4):457–73.

22. Aparicio M, Bellizzi V, Chauveau P, Cupisti A, Ecder T, Fouque D, et al. Protein-restricted diets plus keto/amino acids--a valid therapeutic approach for chronic kidney disease patients. J Ren Nutr. 2012 Mar;22(2 Suppl):S1–21.

23. Buckley BM, Williamson DH. Origins of blood acetate in the rat. Biochem J. 1977 Sep;166(3):539–45.

24. Park J, Kim M, Kang SG, Jannasch AH, Cooper B, Patterson J, et al. Short-chain fatty acids induce both effector and regulatory T cells by suppression of histone deacetylases and regulation of the mTOR-S6K pathway. Mucosal Immunol. 2015 Jan;8(1):80–93.

25. Pluznick JL, Protzko RJ, Gevorgyan H, Peterlin Z, Sipos A, Han J, et al. Olfactory receptor responding to gut microbiota-derived signals plays a role in renin secretion and blood pressure regulation. Proc Natl Acad Sci U S A. 2013 Mar;110(11):4410–5.

26. Park J, Goergen CJ, HogenEsch H, Kim CH. Chronically Elevated Levels of Short-Chain Fatty Acids Induce T Cell-Mediated Ureteritis and Hydronephrosis. J Immunol. 2016 Mar;196(5):2388–400.

27. Brown SA, Brown CA, Crowell WA, Barsanti JA, Allen T, Cowell C, et al. Beneficial effects of chronic administration of dietary omega-3 polyunsaturated fatty acids in dogs with renal insufficiency. J Lab Clin Med. 1998 May;131(5):447–55.

28. Hall JA, Yerramilli M, Obare E, Yerramilli M, Panickar KS, Bobe G, et al. Nutritional Interventions that Slow the Age-Associated Decline in Renal Function in a Canine Geriatric Model for Elderly Humans. J Nutr Health Aging. 2016;20(10):1010–23.

29. Li R, Dai J, Kang H. The construction of a panel of serum amino acids for the identification of early chronic kidney disease patients. J Clin Lab Anal. 2018 Mar;32(3).

30. Duranton F, Lundin U, Gayrard N, Mischak H, Aparicio M, Mourad G, et al. Plasma and urinary amino acid metabolomic profiling in patients with different levels of kidney function. Clin J Am Soc Nephrol. 2014 Jan;9(1):37–45.

31. Bauer JE. Lipoprotein-mediated transport of dietary and synthesized lipids and lipid abnormalities of dogs and cats. J Am Vet Med Assoc. 2004 Mar;224(5):668–75.

32. Behling-Kelly E. Serum lipoprotein changes in dogs with renal disease. J Vet Intern Med. 2014;28(6):1692–8.

33. Thrall MA, Weiser G, Allison RW, Campbell TW. Veterinary Hematology and Clinical Chemistry. 2nd ed. Wiley-Blackwell; 2012.

34. Xenoulis PG, Steiner JM. Canine hyperlipidaemia. J Small Anim Pract. 2015 Oct;56(10):595–605.

